# Deconvolution of omics data in Python with Deconomix – cellular compositions, cell-type-specific gene regulation, and background contributions

**DOI:** 10.1101/2024.11.28.625894

**Authors:** Malte Mensching-Buhr, Thomas Sterr, Dennis Völkl, Nicole Seifert, Jana Tauschke, Laurenz Engel, Austin Rayford, Oddbjørn Straume, Sushma Nagaraja Grellscheid, Tim Beissbarth, Helena U. Zacharias, Franziska Görtler, Michael Altenbuchinger

## Abstract

**Background:** Gene expression profiles derived from heterogeneous bulk samples contain signals from various cell populations. Cell-type deconvolution approaches are computational tools to reverse engineer the composition of bulks in term of cell populations. Accurate estimates of cell compositions are crucial for identifying cell populations relevant for disease. Moreover, analyses, such as the identification of differentially expressed genes, can be confounded by cellular composition, as differences in gene expression may arise from both variations in cellular composition and gene regulation.

**Results:** We present Deconvolution of omics data (Deconomix) – a comprehensive toolbox for the cell-type deconvolution of bulk transcriptomics data, available as a Python package and standalone graphical user interface. Deconomix stands apart from competing solutions with rich functionality and highly efficient implementations. It facilitates (A) the inference of cellular compositions from bulk transcriptomics data, (B) the machine learning-based optimization of gene weights to resolve small cell populations and to disentangle phenotypically related cells, (C) the inference of background contributions which otherwise would deteriorate cell-type deconvolution, and (D) population estimates of cell-type specific gene regulation. To showcase the application of Deconomix, we present a case study on breast cancer data from TCGA, highlighting subtype-specific cellular compositions and cell-type-specific gene-regulatory programs.

**Conclusion:** We present Deconomix, a comprehensive Python package including a graphical user interface for the inference of cellular compositions, cell-type-specific gene regulation, and background contributions from bulk transcriptomics data.

Key Points

- Deconomix optimizes gene weights to disentangle small cell populations and phenotypically related cells.
- Deconomix estimates cell compositions, unknown background contributions and cell-type specific gene regulation from bulk transcriptomics data.
- A Python package is available as code repository and installable via PyPI [1].
- A standalone graphical user interface is available as code repository [2].
- An exemplary analysis of a breast cancer case study is provided as tutorial [3].

## Introduction

Heterogeneous bulk transcriptomics samples are composed of molecularly diverse cellular populations – the different cell types. Variations in cell type composition in diseases like cancer have been repeatedly observed and their association with clinical or phenotypical data can drive the identification of therapeutic targets [4, 5]. Moreover, typical bulk analysis can be confounded by cellular composition [6]; for instance, in differential gene expression analysis, gene expression differences may arise from variations in both cellular composition and gene regulation. Cell-type deconvolution is a computational strategy to infer cellular compositions from bulk or spatial omics data. Several approaches have been proposed in recent years such as BayesPrism, CIBERSORTx, Scaden, and ADTD [7, 8, 9, 10]. Moreover, comprehensive benchmark studies were conducted, highlighting performance and functional differences between methods [11, 12, 13].

Numerous issues complicate cell-type deconvolution and those are inextricably linked. First, the inference of small cell populations is challenging. However, lowly abundant cell types can be highly relevant to answer disease-related questions. For instance, rare immune cells can be key components in cancer defense [14]. As minor cell populations contribute little to the observed bulk profiles, cell-type deconvolution typically assigns a small cellular proportion to them. However, to establish meaningful associations between cellular abundance and phenotype, class, or patient outcomes, the estimated cellular proportions must also correlate closely with the true proportions. Thus, precise estimates are essential, as coarse estimates can blur association analyses. Second, molecularly related cell types can be hardly disentangled by cell-type deconvolution [15]. This difficulty arises when bulk profiles can be equally well reconstructed by different linear combinations of related cell types. As a consequence, the estimated weights corresponding to cellular proportions become imprecise. Third, if specific cell types are missing in the reference profiles, model results can be compromised [11, 16]. Consequently, all cell types contained in the bulk samples should ideally be also included in the reference matrix [11]. Moreover, if this requirement is not fulfilled, estimated proportions of the remaining cell types will be artificially increased and thus biased [7]. A fourth, related problem is the experimental source of reference profiles. It is widely recognized that cells’ molecular phenotypes are influenced by their environment, which also affects results from cell-type deconvolution [17, 18]. To deal with this issue, it was suggested to select references from niches that are comparable to the bulk samples being studied [17]. In summary, practical needs motivate optimized solutions for cell-type deconvolution and the inference is challenged by domain differences such as hidden background contributions and environmental factors. Moreover, recent methodological successes have substantially broadened the scope of cell-type deconvolution [7, 19], and provide estimates of cell-type specific gene regulation from bulk transcriptomics data. One should note that the described obstacles are strongly connected and cannot be solved in isolation; if cell types are missing in the reference profiles, the inference of the remaining, modeled cell types are biased and, vice versa, incorrect estimates of the modeled cell types can deteriorate the inference of background contributions.

Recognizing the fact that the aforementioned issues and obstacles are connected, we suggest a comprehensive toolbox – Deconvolution of omics mixtures (Deconomix) – to address them jointly. Deconomix implements a broad collection of functions that provide methods and utilities for constructing complete cell-type deconvolution pipelines in Python. Its functionality can be organized into three modules: module 1 addresses the gene selection and weighting. Using single-cell training data, it optimizes a set of gene weights on artificially generated bulk profiles and establishes a reference expression matrix for the cell types present in the training data. This procedure allows to reliably quantify lowly abundant and molecularly related cell types. Module 2 uses these gene weights to infer both cellular proportions and the cellular background from bulk transcriptomics data. Module 3 extends the latter analysis by inferring cell-type specific gene regulation. Moreover, Deconomix provides rich functionality to assess and visualize deconvolution results.

## Findings

### Deconomix analysis modules

We will first briefly introduce the three modules of Deconomix and the main features they comprise.

#### Gene weight optimization

This module is dedicated to the machine-learning based optimization of cell-type deconvolution using single-cell data. Specifically, Deconomix provides a method to generate pseudo bulks from single-cell data, accompanied by a reference expression matrix for all cell types present in the dataset. The pseudo bulks are artificial mixtures obtained by randomly drawing single cell profiles from the input data and by aggregating their gene expression. Given the reference matrix and the mixtures, Deconomix establishes an optimized cell-type deconvolution via a gene weighting: genes which are informative for cell-type deconvolution receive high weights, while those which are irrelevant receive low weights. To achieve this, Deconomix optimizes the Pearson’s correlation between estimated and ground truth compositions, as suggested in [20]. Further, the cosine similarity as alternative optimization strategy. Figure 1A shows the corresponding optimization paths exemplified for Pearson’s correlation, where the dotted black line represents the average correlation (*y*-axis) versus the number of iterations used for model optimization and the colored lines correspond to the cell types included in the reference matrix. Figure 1B shows the corresponding scatter plot for a rare cell population before (gray) and after optimization (red).

**Figure 1.**
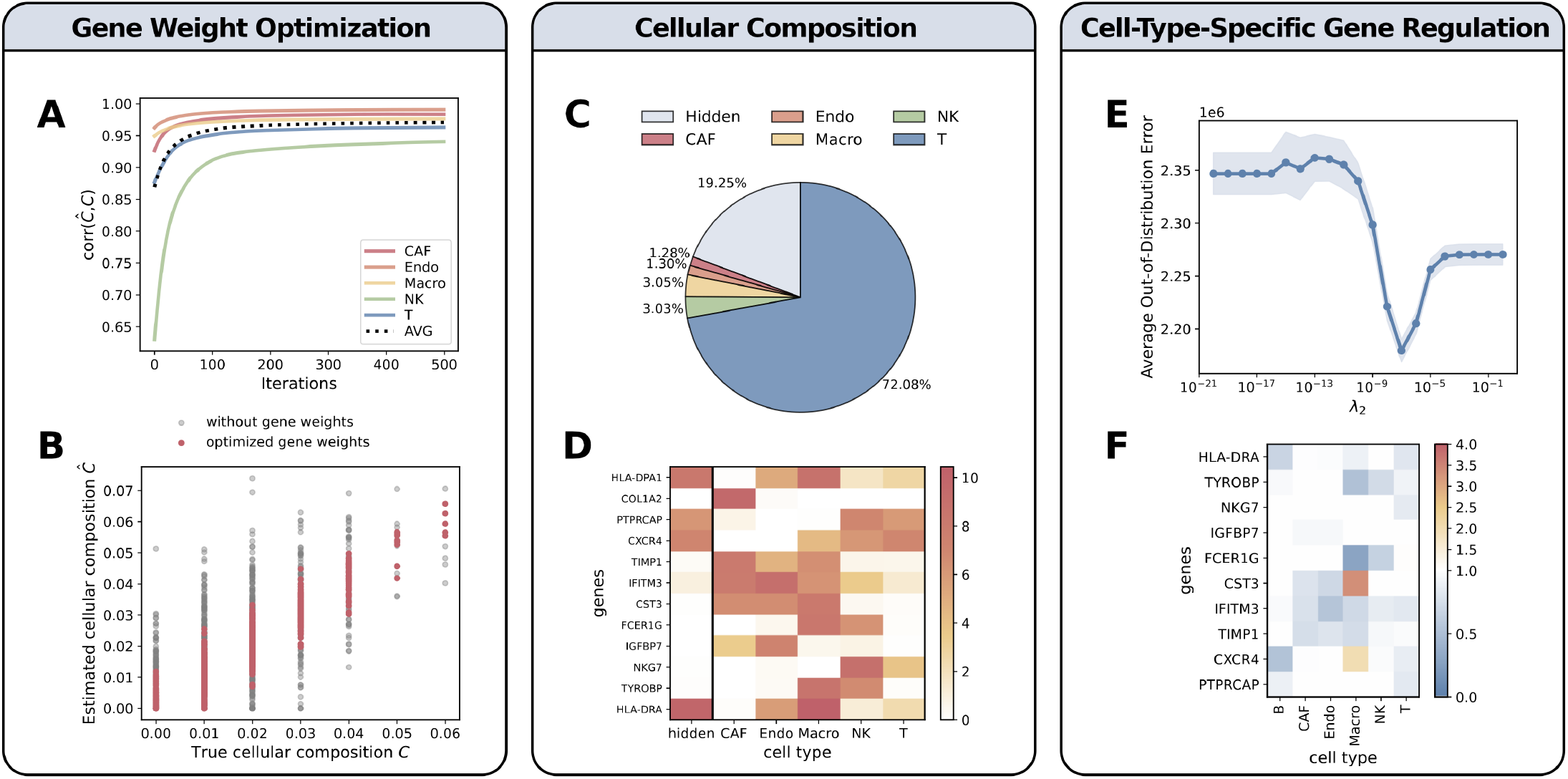
Deconomix Analysis Modules with example realizations: (A) Average and cell-type specific correlation between inferred cellular proportions and respective ground-truth values (*y*-axis) versus the iteration of the Deconomix optimization procedure (*x*-axis). (B) Scatter plot contrasting predictions (*y*-axis) with ground truth values (*x*-axis) before (black) and after Deconomix optimization (red) for a rare cell population (NK cells). (C) Example realization of an inferred cell composition including background contributions. (D) Excerpt of the corresponding reference matrix including an inferred background profile (column “hidden”). (E) Hyper-parameter grid search to infer cell-type specific gene regulation and (F) the corresponding cell-type specific gene regulation matrix.

#### Cellular composition

In this module, bulk data are analyzed utilizing the optimized gene weights and the reference matrix from the first module. It performs both cell-type deconvolution to estimate cellular proportions and infers potential hidden background contributions (referred to as Deconomix+h, Figure 1C and 1D). Estimates of background contributions consist of a consensus background profile across all bulks and the respective bulk-specific proportions. These estimates are optimized via quadratic programming, as proposed in [10] and described in the methods section. The analysis functions are accompanied by functions for model visualization and validation.

#### Cell-type-specific gene regulation

This module is designed to infer cell-type-specific gene regulation along potential background contributions (referred to as Deconomix+h,r). Specifically, it disentangles whether differences in gene expression result from variations in cellular compositions or gene regulation. It requires gene weights and reference profiles established in the first module, and bulk profiles to be analyzed as input. Then, cell-type specific gene regulation is estimated relative to the reference profiles. The algorithmic strategy is outlined in [10] and in the Methods section. This module further performs a hyperparameter search to control overfitting (Figure 1E). The user can select the best hyper-parameter between two different choices for subsequent analysis – a liberal choice, the minimum of the curve, and a more conservative choice, the one standard error rule, described in the methods section. Finally, the Deconomix+h,r model, applied with the chosen hyperparameter, returns the cellular compositions for known cell types and the hidden background with the hidden consensus profile. A matrix of gene-regulation factors Δ_*jk*_ for gene *j* in cell type *k* (Figure 1F) is returned, which can be related to cell-type specific gene expression changes: a factor Δ_*jk*_ > 1 indicates an expression increase in gene *j* in cell type *k* in the bulks relative to the reference profile for cell type *k*.

### Deconomix graphical user interface

Besides the Python package, Deconomix comes with its own graphical user interface (GUI) version. In addition to standalone software for Linux and MacOS, we provide the source code [2] to enable hosting the application on a server. The GUI is designed for non-expert users who want to apply our models to their data without requiring programming expertise.

#### Workflow

Figure 2 shows parts of the GUI and their functionality. The first step is the data import via the input mask shown in Figure 2A. This can be done either via the widely used anndata format (.h5ad) or one can use the file converter tool to generate (.dcx) files, an efficient file format developed for the Deconomix GUI. After importing reference profiles, training and test data, one can start the analysis by optimizing gene weights. In the interface shown in Figure 2C, the user can specify all adjustable parameters, such as number of iterations, number of cells in the artificial bulk mixtures, and number of mixtures. The application then produces pie charts with the cell type distribution for every mixture inferred by a simple Deconomix model, and an interactive heatmap of the top *n* gene weights, where *n* can be specified by the user in the plot panel. A third plot provides information about the correlation between estimated cell type distribution and the simulated ground truth via a cell-type-specific scatter plot. Utilizing these learned gene weights, one can establish more complex Deconomix models, resolving possible hidden cell populations and gene regulation patterns. The Deconomix and the Deconomix+h model can be applied directly. For the Deconomix+h,r model, a hyperparameter can be specified (Figure 2D), or the built-in hyperparameter search can be used. After running the models, the software provides visualizations for the results, including a pie chart for the cellular composition of each analyzed bulk sample. For the cell-type specific gene regulation factors inferred with the Deconomix+h,r model, an interactive heatmap is generated that allows visualization for a user-specified gene set. All plots and results can be exported and used for downstream analyses.

**Figure 2.**
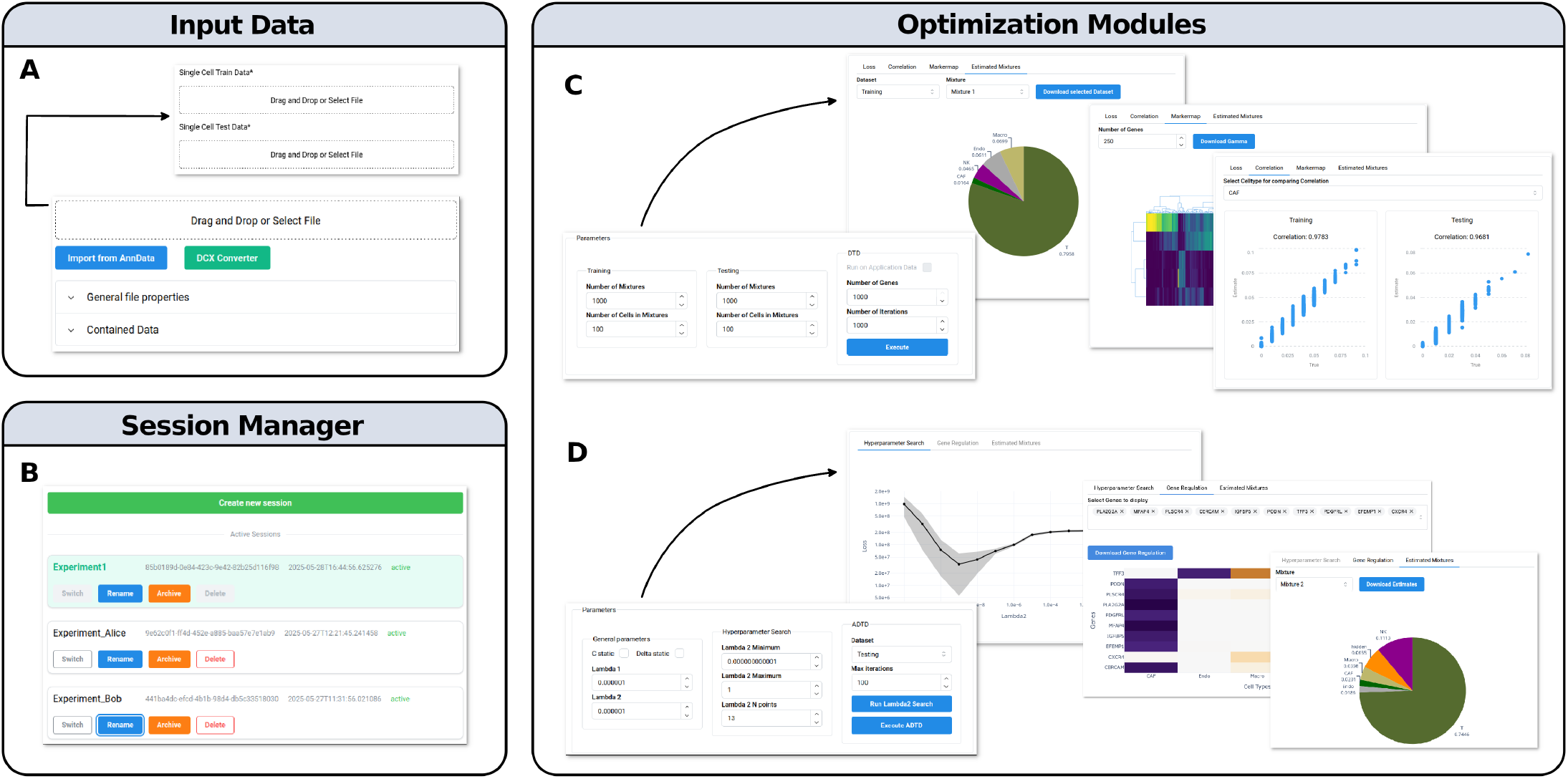
Deconomix Graphical User Interface: The application is locally hosted and enables the users to work with sensitive data. Through the session manager (B) multiple users can work within the same interface. After integrating the data through the interface via .h5ad or .dcx (internal file format; conversion tool can be found on the website) (A) the user can run different Deconomix models (C,D) and the exportable results are visualized with interactive plots, including correlation plots to ground truth values, pie charts of estimated cellular compositions, heatmaps for reference and learned expression profiles, and heatmaps for cell-type-specific gene regulation factors.

#### Session Manager

Upon starting the GUI, the user can either choose to remain in the automatically created session or decide to start a separate working environment via the built-in session manager, shown in Figure 2B. The session manager is especially useful when multiple experiments need to be analyzed in a structured way. But its full potential is shown, when the Deconomix GUI is hosted on a server. The GUI is then running on a specified port (default 8050) and can be accessed through any browser in the local system. The application can then be used by multiple people, administrated by the session manager. This has several advantages. The GUI version can be hosted on a high performance computing cluster to benefit from increased computational resources in terms of RAM and CPU cores. Further, the GUI can be hosted in a secure server environment to handle sensitive data. Then, the data flow is limited to the local network. An additional benefit is the ease of maintenance, as soon as updates for Deconomix are available, only one instance of the GUI version needs to be updated.

### Breast cancer use case

Next, we demonstrate the capabilities of Deconomix in a breast cancer case study, exploring three different settings. The first two settings use single-cell data of breast cancer specimens downloaded from the DISCO database [21, 22]. Those were used to establish artificial bulks for both model training and validation. Both scenarios have the advantage that a clear ground truth is available and that models can be validated accordingly. In contrast, scenario three addresses a real world application on breast cancer bulks from TCGA ([23, 24]), where the single-cell data from DISCO are used for model establishment only. In this scenario, no rigorous validation is feasible, since no established ground truth is available. However, several biological plausibility checks will be provided.

Furthermore, one should note that Deconomix is a comprehensive collection of cell-type deconvolution methods benchmarked previously [10, 20], with modules addressing gene selection and weighting, background inference and the estimation of cell-type specific gene expression. Thus, it contains components that can be used independently, jointly, or even in combination with alternative methodology. For this reason, we refer to the original publications for detailed benchmark analyses, and focus on a typical workflow comprising model development, evaluation, and application to a real world scenario. The Deconomix models used in this study are listed in Table 1, together with their algorithmic background and the quantities they can estimate. All relevant data resources and the necessary preprocessing steps are summarized in the methods section.

**Table 1.** Overview of the Deconomix models used in the breast cancer use case study, summarizing the underlying algorithms, the use of gene weight optimization, and the quantities inferred by each model: Cellular contributions of known cell types **C**, cellular contributions of a hidden background **c**, consensus expression profile of the hidden background **x** and cell-type-specific gene regulation factors Δ.

| Model | Algorithm | Gene Weights | Inferred outputs |  |  |  |
| --- | --- | --- | --- | --- | --- | --- |
| | | | C | c | x | $\Delta$ |
| Naive | NMF with constant feature matrix | — | ✓ | — | — | — |
| Deconomix | DTD | ✓ | ✓ | — | — | — |
| Deconomix+h | DTD + ADTD with $\lambda_1 \rightarrow \infty, \lambda_2 \rightarrow \infty$ | ✓ | ✓ | ✓ | ✓ | — |
| Deconomix+h,r | DTD + ADTD with $\lambda_1 \rightarrow \infty, \lambda_2$ via hyperparameter search | ✓ | ✓ | ✓ | ✓ | ✓ |

#### Scenario 1: A controlled simulation

Scenario 1 is a controlled setting, where we retrieved only cancer cases from DISCO together with corresponding cell-type annotations aggregated into minor cell populations (Table S1). We first split the data randomly into a training and a test set with compositions summarized in Table S2. For illustration purposes, we removed all samples labeled as cancer epithelial cells from the training set, while we retained them in the test data to simulate a domain shift in terms of an unknown expression contribution. Using Deconomix functions, we generated 20,000 artificial bulk mixtures on the training set and determined an average expression profile for each cell type contained in the training data to establish a reference matrix. For validation, we generated 10,000 artificial bulks from the test data and split them into ten test folds for validation. Then, we applied Deconomix to (1) perform a naive cell type deconvolution with equally weighted genes and the afore derived reference matrix, (2) optimize gene weights using the artificial training bulks, and (3) infer a potential background contribution (Deconomix+h). Scenario 1 offers a clear ground truth in terms of underlying cellular compositions in the test bulks. Therefore, we evaluated models (1) to (3) on the test data and calculated Spearman’s correlation coefficient ρ between estimated and true cellular proportions with results summarized in Table S3. The results nicely illustrate the performance benefit of each individual component: while the naive model yielded an average ρ of 0.335 across 18 known cell types, Deconomix with optimized gene weights significantly improved this to an average ρ of 0.634. Notably, B cells improved from 0.185 to 0.793. The naive model further struggled with CD4+ and CD8+ T-cells, yielding 0.414 and 0.319, while Deconomix boosted these values to 0.813 and 0.685. We further observed that Deconomix+h further enhanced performance, achieving an average ρ of 0.658 for known cell types and reliably estimating cellular proportions of an unknown cell type with ρ = 0.655. Thus, in summary, we observed that gene-weight optimization improves deconvolution results and that results become more reliable if background contributions are taken into account.

#### Scenario 2: A controlled simulation with substantial domain shifts

Next, we tested Deconomix’s ability to generalize across application domains by training models on DISCO single-cell data from healthy tissue and validating them on DISCO breast cancer data. In this context, we treated cancer epithelial cells as hidden contributions and used the major cell type annotation denoted in Table S1 to reduce model complexity. An overview of the training and test data compositions is provided in Table S4. As previously, we used the Deconomix functions to generate the training and test bulks, and to evaluate the naive model, the Deconomix model, and the Deconomix+h model. The results, shown in Table S5, indicate that Deconomix models with optimized gene weights significantly outperform the naive model, achieving average correlation values of 0.68 compared to 0.37. The Deconomix+h performed similar and additionally estimated the proportions of the background contribution with ρ = 0.564. In summary, despite an underlying domain shift, both Deconomix models performed well, with almost all cell types exceeding ρ = 0.6. Notably, performance was only compromised significantly for healthy epithelial cells with performance ranging from 0.244 (naive model) and 0.262 (Deconomix), to 0.321 (Deconomix+h). This is most likely a result of the present domain shift and the binary classification of epithelial cells into healthy and cancerous, while these lie, in reality, on a gradient between healthy and cancerous.

#### Scenario 3: Deconomix analysis of breast cancer bulks from TCGA

Finally, we applied Deconomix to real bulk expression data from the TCGA-BRCA cohort [23, 24]. These data comprise, in total, *n* = 1, 176 samples from four breast cancer subtypes: Luminal A (*n* = 637), Luminal B (*n* = 233), HER2+ (*n* = 91), and Basal-like (*n* = 215). In line with scenario 2, we established the Deconomix models on the healthy single-cell data. This has the advantage that the estimated cell-type specific gene regulation can be interpreted relative to healthy controls. Furthermore, we mitigated potential platform effects by gene-wise platform adaptation factors, and selected *p* = 5,000 genes present in both the single-cell and bulk data (see Methods). Given the comparatively complex scenario comprising potential background contributions and likely cell-type specific gene regulation between healthy single-cell and the cancerous breast cancer bulks, we applied Deconomix in model “Deconomix+h,r” (estimating hidden background and cell-type specific gene regulation). This model requires calibration of a hyperparameter λ_2_ to control overfitting. Deconomix automatically screens a prespecified series of λ_2_ values and finally selects the value with the lowest cross-validation error or the one according to the one standard error rule (see Methods). For the analyses presented here, we rely on the latter choice, which generally yields the more conservative model. At this point, the user already receives all results of interest: cell composition estimates, background contributions, and cell-type specific gene regulation. There are several possible downstream analysis options. First, cell compositions can be correlated with phenotypes using test strategies. Table 2 summarizes significant cell-composition differences between luminal A, luminal B, HER2+, and basal-like breast cancers obtained via a Kruskal-Wallis test, containing both raw *p*-values and those corrected for multiple testing according to Benjamini and Hochberg [25]. Second, cell-type specific gene regulation can be explored. We visualized the top 20 upregulated and downregulated genes across four cancer subtypes in heatmaps (Figures 3 and S2). Both coherent and subtype specific gene expression patterns were observed, as illustrated for the top 40 upregulated genes per subtype in Venn diagram Figure 4. For instance, Luminal A, Luminal B, and Her2+ share several upregulated genes, while the Basal-like subtype is more unique, with 25 genes not overlapping. Notably, five genes – B2M, IFITM1, TXN, IGHG1, and HMOX1 – are consistently upregulated across all subtypes. B2M, linked to immune regulation, is predicted to be underexpressed in epithelial cells, possibly contributing to immune escape [26]. IGHG1 is upregulated in B-cells across subtypes, in line with adaptive immune activation [27]. Third, Deconomix includes functions to perform enrichment analyses with respect to the estimated cell-type specific gene expression estimates, where both Fisher’s exact test and the Mann-Whitney U-test can be used. This analysis can be performed for the individual breast cancer subtypes, and can be tailored to specific cell types of interest. For illustration, we conducted a GSEA using GSEApy [28], focusing on the MSigDB C7 database of immunologic gene sets [29, 30]. The top 15 hits, presented in Table S6, show significant enrichment in T-cell activation, regulatory T-cell signals, and innate immunity signatures, confirming the anticipated immune response in diseased tissue.

**Table 2.**
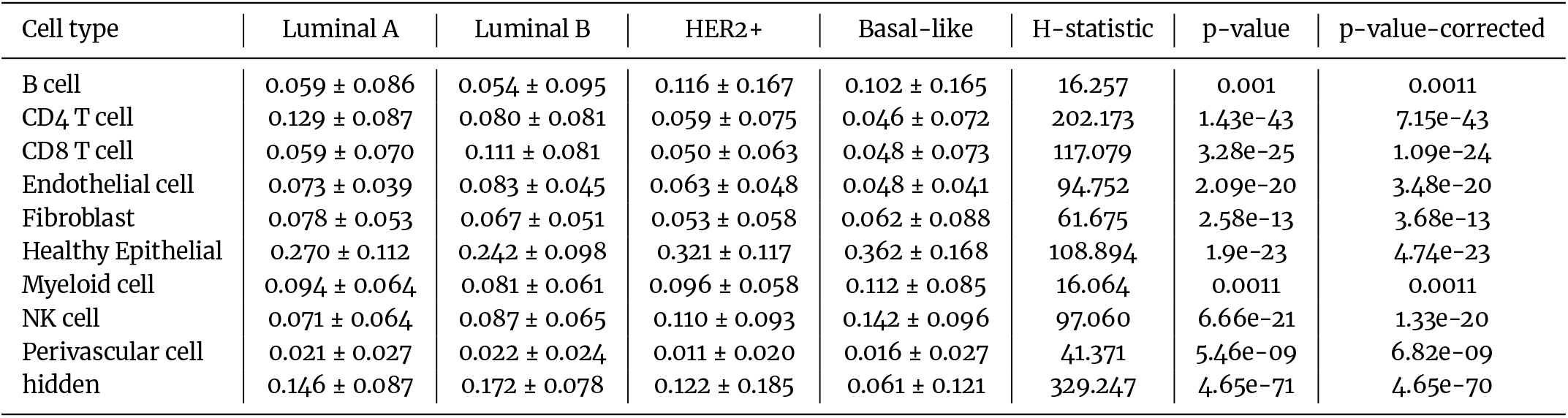
Average cellular composition of four breast cancer subtypes within the TCGA-BRCA cohort [23, 24] estimated by a Deconomix+h,r model which infers hidden background contributions and cell-type specific gene regulation. Values denoted as mean± one standard deviation. H-statistic and p-values determined by Kruskal-Wallis tests to identify significant differences in the composition of the different breast cancer subtypes. P-values adjusted for multiple testing by controlling the false-discovery rate (Benjamini-Hochberg [25]).

**Figure 3.**
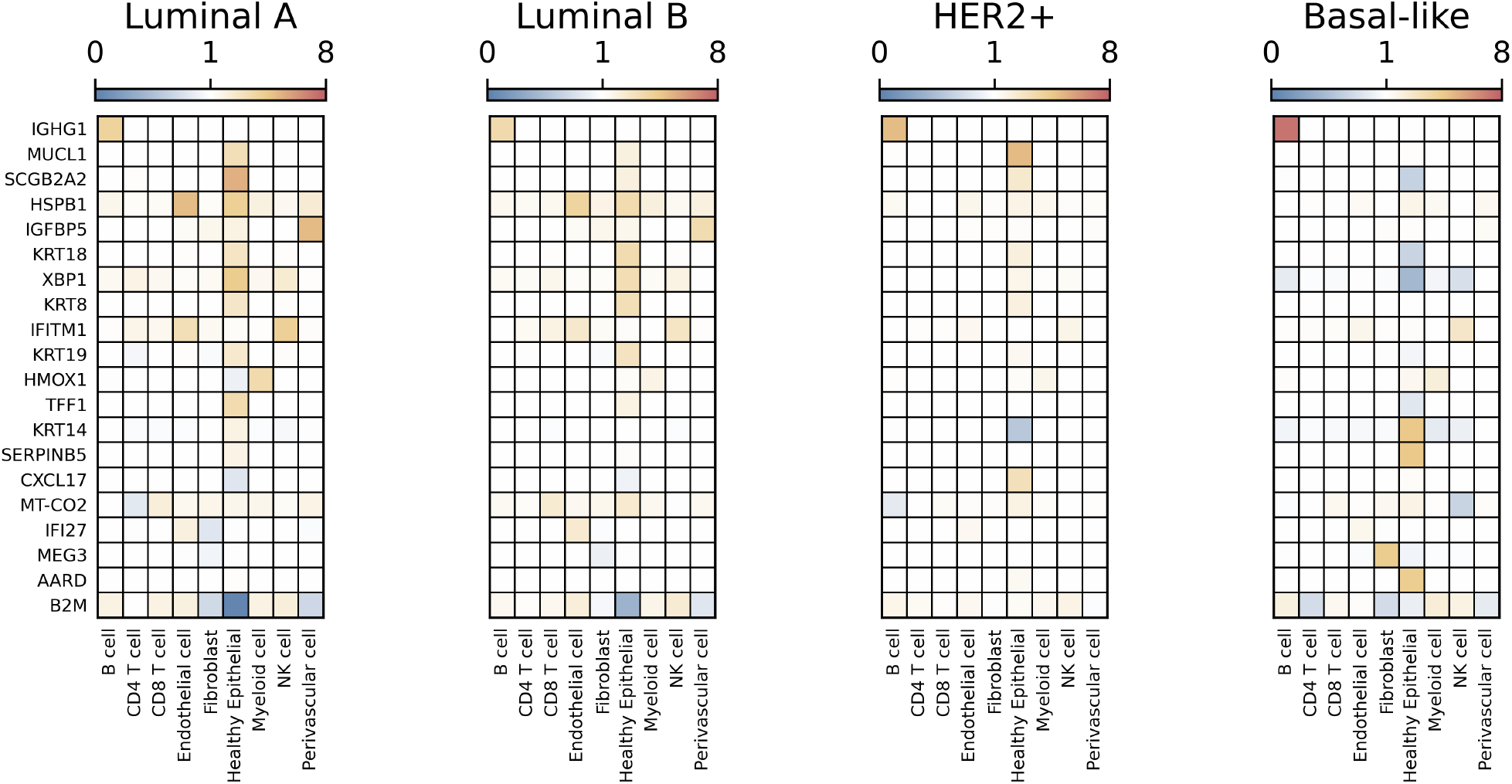
Heatmap of the 20 genes most upregulated across the four breast cancer subtypes. Values between 0 and 1 (blue) correspond to down-regulation, white to no regulation, values above 1 (yellow over orange and red) correspond to positive gene regulation.

**Figure 4.**
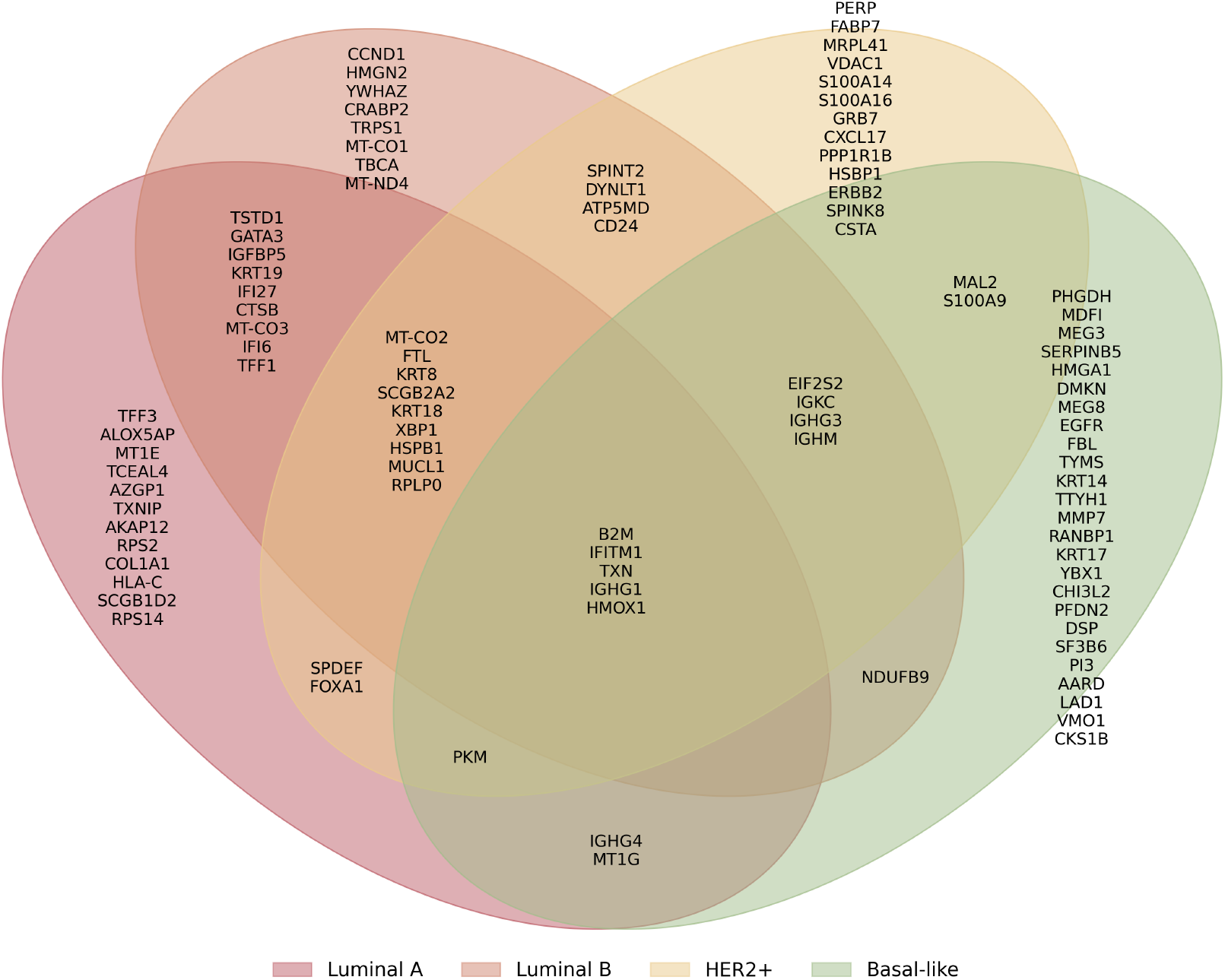
Venn diagram illustrating the overlap of the 40 top up-regulated genes in any cell type across the four breast cancer subtypes within TCGA breast cancer data [23, 24], inferred by a Deconomix+h,r model with λ_2_ = 1e-11.

## Discussion

Computational approaches for cell-type deconvolution have evolved substantially in recent years, starting from naive strategies based on ordinary least squares to complex models which infer not only cellular proportions, but also potential background contributions and cell-type specific gene regulation [7, 10, 19, 31, 32, 33]. The implementation of such complex models requires dedicated algorithmic solutions that typically also require more computing resources. Moreover, advanced methods are usually more difficult to apply and require substantial programming skills. In recent years, we established several cell-type deconvolution approaches, where we systematically addressed different analysis bottlenecks [10, 34, 35, 20], such as potential background contributions, gene-regulatory adaptation, and the reliability cell composition estimates. Notably, all these bottlenecks are inevitably connected. Therefore, we combined them in Deconomix – a user-friendly Python pipeline and stand-alone GUI – for users with no or little programming experience. With Deconomix, users can rely on a highly efficient state-or-the-art pipeline for cell composition analyses that accounts for dominant analysis pitfalls. Moreover, Deconomix has rich functionality, such as functions for generating pseudo-bulk data from single-cell datasets, to perform, visualize and benchmark cell-type deconvolution, conducting hyperparameter searches for advanced models, and to conduct gene-set enrichment analysis to explore cell-type specific gene regulation.

Comprehensive software suites for cell-composition analysis are scarce and alternatives are highly needed. For instance, Omnideconv stands out as a unified framework for utilizing and benchmarking single-cell-informed cell-type deconvolution methods [36]. It currently supports 12 different approaches. In the same spirit, Decon-Benchmark offers utilities for comprehensive benchmark analysis [37]. However, in contrast to Deconomix, both lack a coherent workflow to systematically guide users through outlined challenges and analytical steps in cell-type deconvolution analysis.

In summary, Deconomix is a comprehensive toolbox for cell-type deconvolution, providing state-of-the-art algorithmic solutions for the inference of cellular proportions, background contributions, and cell-type specific gene regulation. It is further designed for users with little programming knowledge and can also be applied via a user-friendly GUI.

## Methods

### Nomenclature

Let 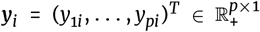 represent the bulk RNA profile of sample *i* containing gene expression levels for *p* genes on a natural scale (not on log scale). Further, let 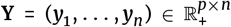 be a matrix containing *n* bulk profiles in its columns. The archetypical cell-type reference profiles are denoted as 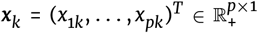, where *k* = 1, …, *q* and 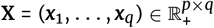 is the reference matrix containing the *q* reference profiles in its columns. We further use the nomenclature A _*k*, ·_ to indicate the *k*th row of an arbitrary matrix **A**.

### Gene Weight Optimization

The key idea behind cell-type deconvolution is to express the observed bulk profiles as a linear combination of reference profiles, ***y***_*i*_ = **X*c***_*i*_ + **ϵ**_*i*_, where ***c***_*i*_ = (*c*_*i*1_, …, *c*_*iq*_)^*T*^ are the regression weights for the *q* cell types in sample *i* and **ϵ**_*i*_ = (ϵ_1*i*_, …, ϵ_*pi*_) the corresponding gene residuals. For a set of bulk profiles, this equation becomes

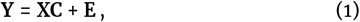

where 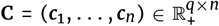 contains the regression weights for *n* samples in its columns. The corresponding model residuals are collected in E = (ϵ_1_,… ϵ_*n*_) ∈ ℝ^*p*×*n*^. The regression weights can be interpreted as cellular compositions under normalization constraints; if each column of **Y** and **X** sums up to one and assuming that there are no missing reference profiles in **X**, one obtains approximately ∑_*k*_ *c*_*ik*_ = 1 for *i* = 1, …, *n* [18, 10]. For computational efficiency, we relied on non-negative least squares,

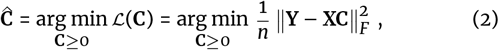

which can also be minimized for each ***c***_*i*_ individually. The key step of loss-function learning is to generalize the loss to

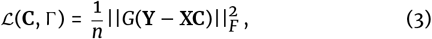

with *G* = diag(*g*_1_, …, *g*_*p*_) ∈ ℝ^*p×p*^ containing weights *g*_*j*_ for each gene *j*. Then, for each gene *j*, the corresponding term in the sum of squared residuals is weighted by a factor 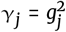. Thus, to improve estimates of cellular compositions, genes can be weighted according to their importance for cell-type deconvolution; genes which improve cell-type deconvolution are up-weighted while those which do not improve cell-type deconvolution are down-weighted. This is achieved by optimizing an outer loss function, for which we provide two different choices. In line with the Digital Tissue Deconvolution algorithm (DTD) [20], we implemented the outer objective function

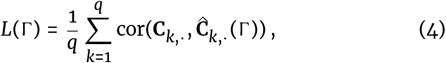

which is maximized with respect to Γ = diag(γ_1_, …, γ_*p*_) subject to 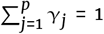, given a reference matrix **X** and training mixtures **Y** of known cellular composition **C**. To accelerate optimization, 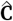 is determined from ordinary least squares, where all values *c*_*ik*_ ≤ 0 are set to zero. The training mixtures can be generated, e.g., using single-cell data [20].

One should note that 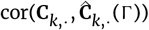 is the correlation between ground truth proportions and their estimates for cell type *k*. Consequently, each modeled cell type is equally important for the outer loss, which is crucial for the correct inference of minor and molecularly related cell populations. As an alternative choice for the outer loss function, we implemented the cosine similarity given by

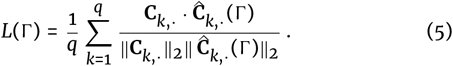

For implementation of both outer loss functions, the Pytorch machine learning library (version 2.3.1) was used [38]. Respective functions for preparing training data and the gene-weight optimization itself are provided by Deconomix.

### Inferring background contributions and cell-type-specific gene regulation factors

The algorithm Adaptive Digital Tissue Deconvolution (ADTD) [10] expands the introduced supervised loss-function learning routine with an unsupervised approach to refine deconvolution results. It utilizes the gene weights Γ determined as outlined above, but additionally accounts for potential background contributions not contained in the reference matrix and for environmental factors. The ADTD loss is defined as

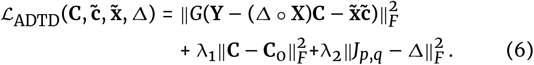

The corresponding optimization problem is

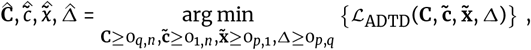

subject to 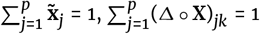 for all *k*, and 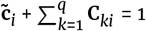. Here, the term 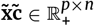 models background contributions, where the unknown background 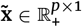 contributes with proportions 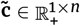 for the *n* bulks. Further, Δ ° **X** represents an adapted reference matrix, where 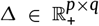 contains gene and cell-type specific rescaling factors which are multiplied element-wise with the entries of **X** via the Hadamard product °. The re-scaling factors Δ_*jk*_ can be interpreted as cell-type-specific gene regulation; a coefficient of Δ_*jk*_ = 2 corresponds to a two-fold increase, Δ_*jk*_ = 1 to no gene regulation, and Δ_*jk*_ = 0.5 to a halved expression in gene *j* and cell type *k* relative to the reference, respectively. Eq. (6) further contains two regularization terms. The term 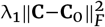regularizes the cellular proportions **C** and biases their estimates to pre-specified proportions **C**_0_. Deconomix determines **C**_0_ by minimizing the ADTD loss with Δ_*jk*_ = 1 for all *j, k*. Thus **C**_0_ corresponds to the ADTD solution without environmental adaptation. The solution 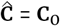 is retrieved for λ_1_→ ∞ . The second regularization term 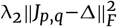 biases the environmental adaptation factors towards the solution without adaptation (Δ_*jk*_ = 1 for all *j, k*).

### Hyperparameter tuning

The Deconomix model with a hidden background (Deconomix+h) assumes that the reference profiles do not require adaptation (Δ_*jk*_ = 1 for all *j, k*). Thus, the user directly obtains the ADTD solution for λ_1_→ ∞, λ_2_ → ∞. Despite its simplicity, this solution already provides a remarkably accurate estimate of cellular proportions **C**, the background profile 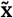 and respective background proportions 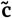 [10].

A more complex Deconomix model with a hidden background and cell-type-specific gene regulation (Deconomix+h,r) is intended to infer population estimates of cell-type-specific gene regulation alongside with the outputs described for the other models. These estimates are encoded in the adaptation factors Δ. In order to provide a computationally cheap strategy to infer Δ without extensive hyperparameter screening, we used the following strategy:

i. Set **C** to the solution without cellular adaptation (**C** = **C**_0_).
ii. Set the gene weights to

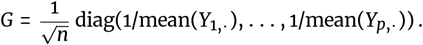

This choice ensures that genes contribute comparably to the optimization problem and that a single λ_2_ value can be used for different classes.
iii. Split the data into batches. When Deconomix is used with standard settings, the data are split in five batches of approximately equal size.
iv. Use *k* – 1 of the *k* data batches to learn an ADTD model for a series of λ_2_ values and record the loss on the left-out batch. Repeat this procedure such that every batch served for model validation once (*k*-fold cross validation).

The former procedure provides an estimate of the generalization error together with error intervals. Deconomix provides two choices to select λ_2_. The first choice is named “lambda2.min” and returns the λ_2_ value with the lowest generalization error. The second choice is more conservative and is named “lambda2.1se”. It is defined as the minimum value that λ_2_ can take such that it is still outside the one standard error interval obtained for λ_2_ = lambda.min, in line with [39]. The latter choice was used for the presented use case analysis.

### Breast cancer use case and data preprocessing

We retrieved single-cell gene expression data from healthy and cancerous breast tissue from the DISCO database [21, 22]. The obtained samples contained 34 fine-grained cell-type labels, which we aggregated into broader cell-type classes informed by known cell lineages. We mapped the original labels to a set of major cell types, featuring nine different classes, and a set of minor cell types featuring 19 classes, shown in Supplementary Table S1. We computed average expression profiles for each originally labeled cell type, which we concatenated into a reference expression matrix, comprising values for all genes and each original cell-type label. Genes were then ranked decreasingly by variance between cell types. TCGA bulk expression data [23, 24] featuring breast cancer bulk expression samples were imported and restricted to genes available in the variance-ranked list. Then, the bulk data were normalized to 10,000 counts per samples, and used to compute gene-wise mean expression values µ_TCGA_. Simultaneously, DISCO cancer single-cell samples were selected by restricting to breast cancer samples derived from solid tumors only and labeling cells according to the minor cell type annotations denoted in Table S1. Genes were selected according to the same variance-based ordering and the data were normalized to 10,000 counts per sample. Artificial bulk mixtures were simulated from these samples, renormalized to 10,000 counts per mixture, and used to compute per-gene mean expression values µ_DISCO_.

To account for platform-specific differences between TCGA and DISCO, we estimated gene-wise conversion factors α:

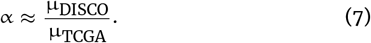

In absence of platform-specific effects, we expect that each gene has a similar mean expression across the two datasets. Strong deviations from α ≈ 1 primarily reflect technical rather than biological differences. Genes for which this similarity assumption was violated, were conservatively filtered by requiring 0.1 ≤ α ≤ 10.0, corresponding to a maximum plausible tenfold change between platforms. From the remaining genes, still ordered by variance across DISCO cell types, we retained the top 5,000 to define the final gene set.

Single-cell samples labeled as cancer epithelial cells under the major annotation, despite originating from healthy samples, were removed from the healthy subset of the DISCO data. After selecting the consensus genes and normalizing to 10,000 counts per sample, we obtained a DISCO cancer and a DISCO healthy dataset.

TCGA bulk samples were processed using the same gene set, adjusted by the estimated conversion factors, and renormalized to 10,000 counts per sample to obtain the final TCGA dataset, which represent the bulk samples to be analyzed in this case study.

## Availability of source code and requirements

- Project name: Deconomix
- Project home page: https://gitlab.gwdg.de/MedBioinf/MedicalDataScience/Deconomix
- Package source code: https://gitlab.gwdg.de/MedBioinf/MedicalDataScience/Deconomix/Deconomix
- Package documentation: https://medbioinf.pages.gwdg.de/MedicalDataScience/Deconomix/Deconomix/
- Breast cancer use case study: https://gitlab.gwdg.de/MedBioinf/MedicalDataScience/Deconomix/Deconomix-case-study
- Graphical user interface: https://gitlab.gwdg.de/MedBioinf/MedicalDataScience/Deconomix/Deconomix-gui
- Programming language: Python >=3.11
- Licence: GPLv3

## Data Availability

All datasets used in the breast cancer case study are publicly available: For the DISCO single-cell human breast atlas we retrieved the file disco_breast_v01.h5ad from the corresponding data repository [22]. Breast cancer bulk expression data from the TCGA-BRCA project [23] were retrieved from the GDC portal [24] via API calls available as script in the code repository of our breast cancer use case study [3]. For the gene set enrichment analysis we retrieved the ImmuneSigDB subset of C7 [29, 30] from the GSEA website as gmt-file c7.immunesigdb.v2025.1.Hs.symbols.gmt from following page: https://www.gsea-msigdb.org/gsea/msigdb/human/collections.jsp

## Declarations

### Authors’ contributions

**Malte Mensching-Buhr:** Software (lead), Methodology (supporting), Validation (supporting), Writing - review and editing (equal). **Thomas Sterr:** Methodology (lead), Software (supporting), Validation (supporting), Writing - review and editing (equal). **Dennis Völkl:** Validation (supporting), Software (supporting), Writing - review and editing (equal). **Nicole Seifert:** Writing - review and editing (equal). **Jana Tauschke:** Validation (supporting). Writing - review and editing (equal). **Laurenz Engel:** Writing - review and editing (equal). **Austin Rayford:** Validation (supporting), Writing - review and editing (equal). **Oddbjørn Straume:** Resources (equal), Writing - review and editing (equal). **Sushma N. Grellscheid:** Resources (equal), Writing - review and editing (equal). **Tim Beissbarth:** Resources (equal), Writing - review and editing (equal). **Helena U. Zacharias:** Writing - review and editing (equal). **Franziska Görtler:** Conceptualization (equal), Supervision (equal), Validation (lead), Writing - review and editing (equal). **Michael Altenbuchinger:** Conceptualization (equal), Supervision (equal), Project Administration (lead), Writing - review and editing (equal).

## Acknowledgements

Malte Mensching-Buhr was supported by the Ph.D. program “Genome Science” - International Max Planck Research School.

## Supplementary Material

**Table S1.**
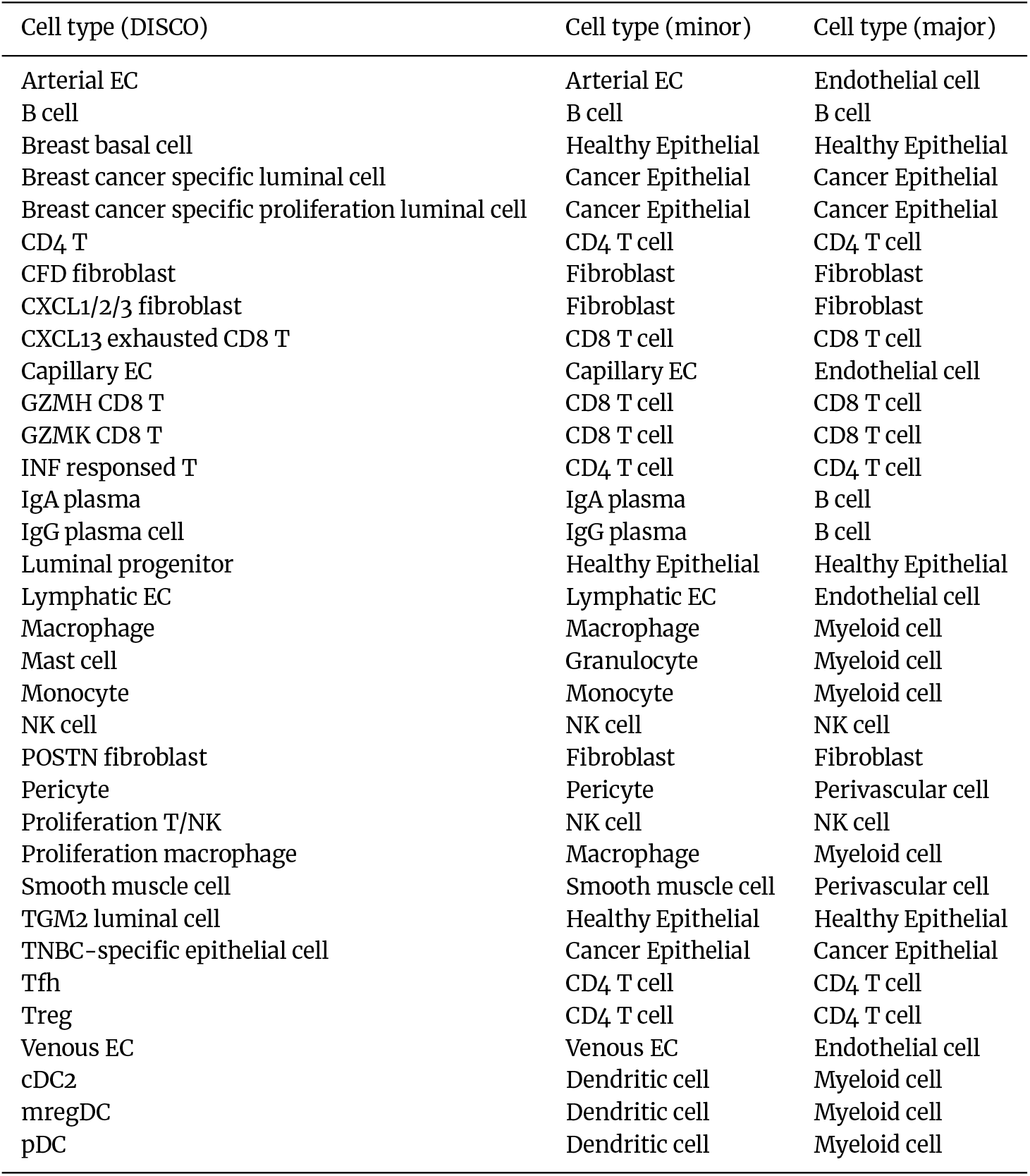
Aggregation of the original, fine-grained cell-type labels from the DISCO single-cell dataset [22] into two sets of more coarse labels. The minor subset features 19, and the major subset nine distinct classes.

**Table S2.** Composition of single-cell samples in a random test/train split of the cancer samples within the DISCO dataset [22]. Relabeled using the minor cell-type labels defined in Table S1. Cancer epithelial cells were dropped from the training set but preserved as ‘hidden contribution’ in the test set.

| Cell type | Train number | Train percentage | Test number | Test percentage |
| --- | --- | --- | --- | --- |
| Arterial EC | 335 | 0.555 | 346 | 0.351 |
| B cell | 2524 | 4.184 | 2505 | 2.538 |
| CD4 T cell | 13376 | 22.175 | 13382 | 13.559 |
| CD8 T cell | 10012 | 16.598 | 9971 | 10.103 |
| Capillary EC | 977 | 1.620 | 1051 | 1.065 |
| Dendritic cell | 1679 | 2.783 | 1571 | 1.592 |
| Fibroblast | 8023 | 13.301 | 7990 | 8.095 |
| Granulocyte | 449 | 0.744 | 439 | 0.445 |
| Healthy Epithelial | 5049 | 8.370 | 5171 | 5.239 |
| IgA plasma | 568 | 0.942 | 570 | 0.578 |
| IgG plasma | 2285 | 3.788 | 2197 | 2.226 |
| Lymphatic EC | 155 | 0.257 | 169 | 0.171 |
| Macrophage | 7968 | 13.210 | 7901 | 8.005 |
| Monocyte | 1112 | 1.844 | 1128 | 1.143 |
| NK cell | 2559 | 4.242 | 2736 | 2.772 |
| Pericyte | 475 | 0.787 | 469 | 0.475 |
| Smooth muscle cell | 1497 | 2.482 | 1477 | 1.496 |
| Venous EC | 1277 | 2.117 | 1276 | 1.293 |
| sum immune cell types | 60320 | 100.000 | 60349 | 61.146 |
| hidden contribution | - | - | 38348 | 38.854 |
| total | 60320 | 100.000 | 98697 | 100.000 |

**Table S3.**
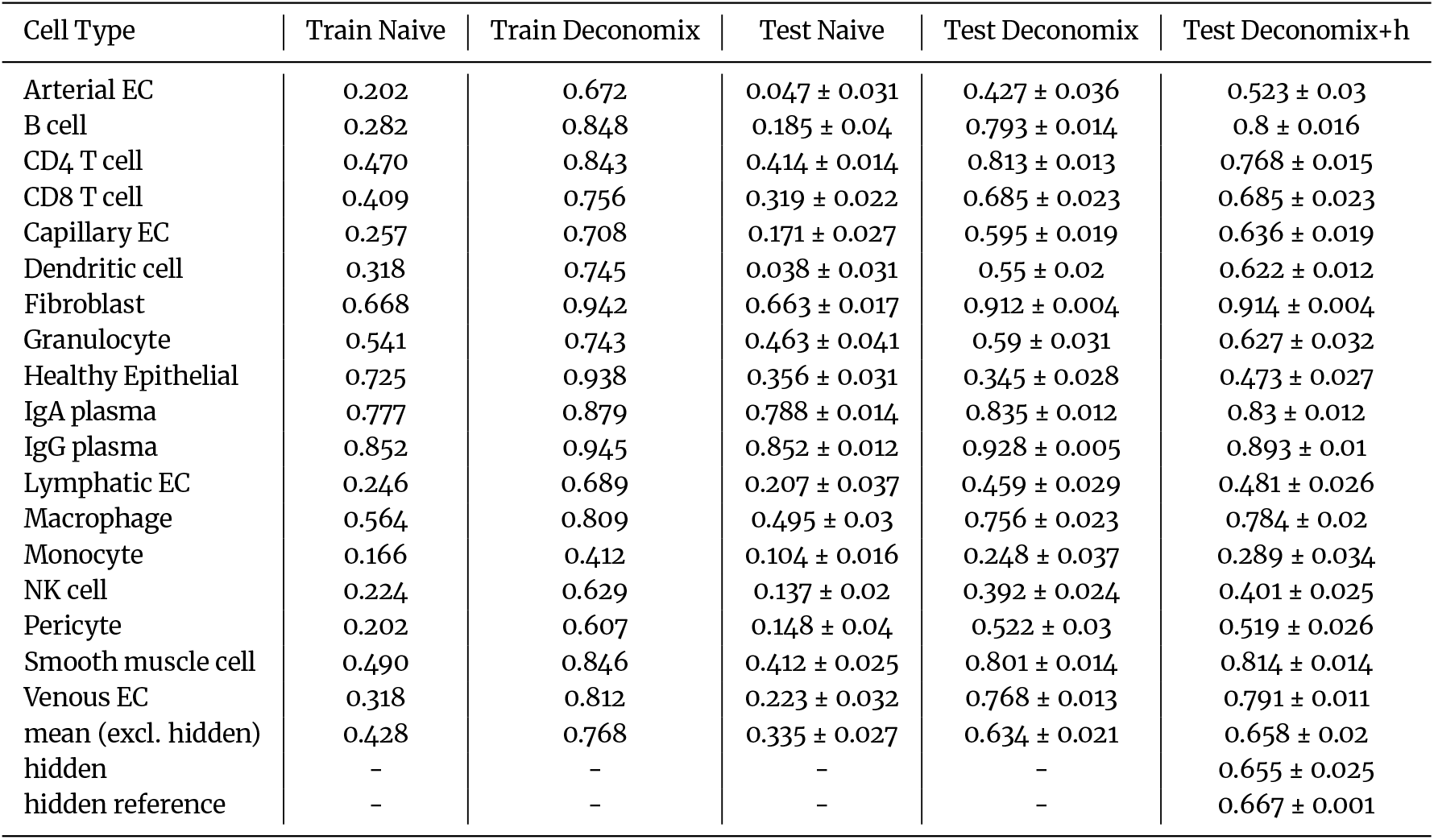
Performance assessment of cell-type deconvolution on bulk profiles simulated from the breast cancer samples within the DISCO dataset[22]. Calculated as Spearman’s correlation coefficient ρ between true and estimated cellular compositions. Performance on training set is shown to illustrate training success. Evaluation on a different test set was conducted on ten folds of the data and is denoted as mean *±*one standard deviation. Three different models are featured: A naive model without Deconomix’ gene weight optimization, a Deconomix model, and a Deconomix+h model, which performs hidden background estimation.

| Cell Type | Train Naive | Train Deconomix | Test Naive | Test Deconomix | Test Deconomix+h |
| --- | --- | --- | --- | --- | --- |
| Arterial EC | 0.202 | 0.672 | 0.047 ± 0.031 | 0.427 ± 0.036 | 0.523 ± 0.03 |
| B cell | 0.282 | 0.848 | 0.185 ± 0.04 | 0.793 ± 0.014 | 0.8 ± 0.016 |
| CD4 T cell | 0.470 | 0.843 | 0.414 ± 0.014 | 0.813 ± 0.013 | 0.768 ± 0.015 |
| CD8 T cell | 0.409 | 0.756 | 0.319 ± 0.022 | 0.685 ± 0.023 | 0.685 ± 0.023 |
| Capillary EC | 0.257 | 0.708 | 0.171 ± 0.027 | 0.595 ± 0.019 | 0.636 ± 0.019 |
| Dendritic cell | 0.318 | 0.745 | 0.038 ± 0.031 | 0.55 ± 0.02 | 0.622 ± 0.012 |
| Fibroblast | 0.668 | 0.942 | 0.663 ± 0.017 | 0.912 ± 0.004 | 0.914 ± 0.004 |
| Granulocyte | 0.541 | 0.743 | 0.463 ± 0.041 | 0.59 ± 0.031 | 0.627 ± 0.032 |
| Healthy Epithelial | 0.725 | 0.938 | 0.356 ± 0.031 | 0.345 ± 0.028 | 0.473 ± 0.027 |
| IgA plasma | 0.777 | 0.879 | 0.788 ± 0.014 | 0.835 ± 0.012 | 0.83 ± 0.012 |
| IgG plasma | 0.852 | 0.945 | 0.852 ± 0.012 | 0.928 ± 0.005 | 0.893 ± 0.01 |
| Lymphatic EC | 0.246 | 0.689 | 0.207 ± 0.037 | 0.459 ± 0.029 | 0.481 ± 0.026 |
| Macrophage | 0.564 | 0.809 | 0.495 ± 0.03 | 0.756 ± 0.023 | 0.784 ± 0.02 |
| Monocyte | 0.166 | 0.412 | 0.104 ± 0.016 | 0.248 ± 0.037 | 0.289 ± 0.034 |
| NK cell | 0.224 | 0.629 | 0.137 ± 0.02 | 0.392 ± 0.024 | 0.401 ± 0.025 |
| Pericyte | 0.202 | 0.607 | 0.148 ± 0.04 | 0.522 ± 0.03 | 0.519 ± 0.026 |
| Smooth muscle cell | 0.490 | 0.846 | 0.412 ± 0.025 | 0.801 ± 0.014 | 0.814 ± 0.014 |
| Venous EC | 0.318 | 0.812 | 0.223 ± 0.032 | 0.768 ± 0.013 | 0.791 ± 0.011 |
| mean (excl. hidden) | 0.428 | 0.768 | 0.335 ± 0.027 | 0.634 ± 0.021 | 0.658 ± 0.02 |
| hidden | - | - | - | - | 0.655 ± 0.025 |
| hidden reference | - | - | - | - | 0.667 ± 0.001 |

**Table S4.** Composition of single-cell samples in a test/train split of healthy samples versus cancer samples within the DISCO dataset [22]. Relabeled using the major cell-type labels defined in Table S1.

| Cell type | Train number | Train percentage | Test number | Test percentage |
| --- | --- | --- | --- | --- |
| B cell | 269 | 0.986 | 10649 | 5.408 |
| CD4 T cell | 3642 | 13.348 | 26758 | 13.588 |
| CD8 T cell | 4792 | 17.563 | 19983 | 10.148 |
| Endothelial cell | 5504 | 20.173 | 5586 | 2.837 |
| Fibroblast | 5311 | 19.466 | 16013 | 8.132 |
| Healthy Epithelial | 1979 | 7.253 | 10220 | 5.190 |
| Myeloid cell | 3564 | 13.063 | 22247 | 11.298 |
| NK cell | 993 | 3.639 | 5295 | 2.689 |
| Perivascular cell | 1230 | 4.508 | 3918 | 1.990 |
| sum immune cell types | 27284 | 100.000 | 120669 | 61.280 |
| hidden contribution | - | - | 76249 | 38.721 |
| total | 27284 | 100.000 | 196918 | 100.000 |

**Table S5.** Performance assessment of different cell-type deconvolution models established on healthy single-cell data and applied bulk profiles simulated from breast cancer samples both within the DISCO dataset [22]. Calculated as Spearman’s correlation coefficient ρ between true and estimated cellular compositions. Performance on training set is shown to illustrate training success. Evaluation on a different test set was conducted on ten folds of the data and is denoted as mean *±* one standard deviation. Three different models are featured: A naive model without Deconomix’ gene weight optimization, a Deconomix model, and a Deconomix+h model, which performs hidden background estimation. The naive model estimated the abundance of CD4 T cells in the test set to be zero which is why no correlation coefficient could be calculated for that field.

| Cell Type | Train Naive | Train Deconomix | Test Naive | Test Deconomix | Test Deconomix+h |
| --- | --- | --- | --- | --- | --- |
| B cell | 0.581 | 0.826 | 0.599 ± 0.026 | 0.87 ± 0.009 | 0.836 ± 0.01 |
| CD4 T cell | 0.245 | 0.808 | - | 0.758 ± 0.013 | 0.752 ± 0.011 |
| CD8 T cell | 0.241 | 0.798 | 0.005 ± 0.024 | 0.617 ± 0.029 | 0.601 ± 0.029 |
| Endothelial cell | 0.753 | 0.960 | 0.341 ± 0.019 | 0.792 ± 0.009 | 0.794 ± 0.011 |
| Fibroblast | 0.785 | 0.970 | 0.670 ± 0.022 | 0.857 ± 0.008 | 0.861 ± 0.01 |
| Healthy Epithelial | 0.733 | 0.942 | 0.244 ± 0.025 | 0.262 ± 0.025 | 0.321 ± 0.03 |
| Myeloid cell | 0.884 | 0.957 | 0.744 ± 0.015 | 0.924 ± 0.003 | 0.919 ± 0.003 |
| NK cell | 0.271 | 0.750 | 0.183 ± 0.036 | 0.398 ± 0.02 | 0.359 ± 0.024 |
| Perivascular cell | 0.680 | 0.912 | 0.196 ± 0.028 | 0.662 ± 0.02 | 0.660 ± 0.019 |
| mean (excl. hidden) | 0.575 | 0.880 | 0.373 ± 0.024 | 0.682 ± 0.015 | 0.678 ± 0.016 |
| hidden | - | - | - | - | 0.564 ± 0.016 |
| hidden reference | - | - | - | - | 0.550 ± 0.001 |

**Table S6.**
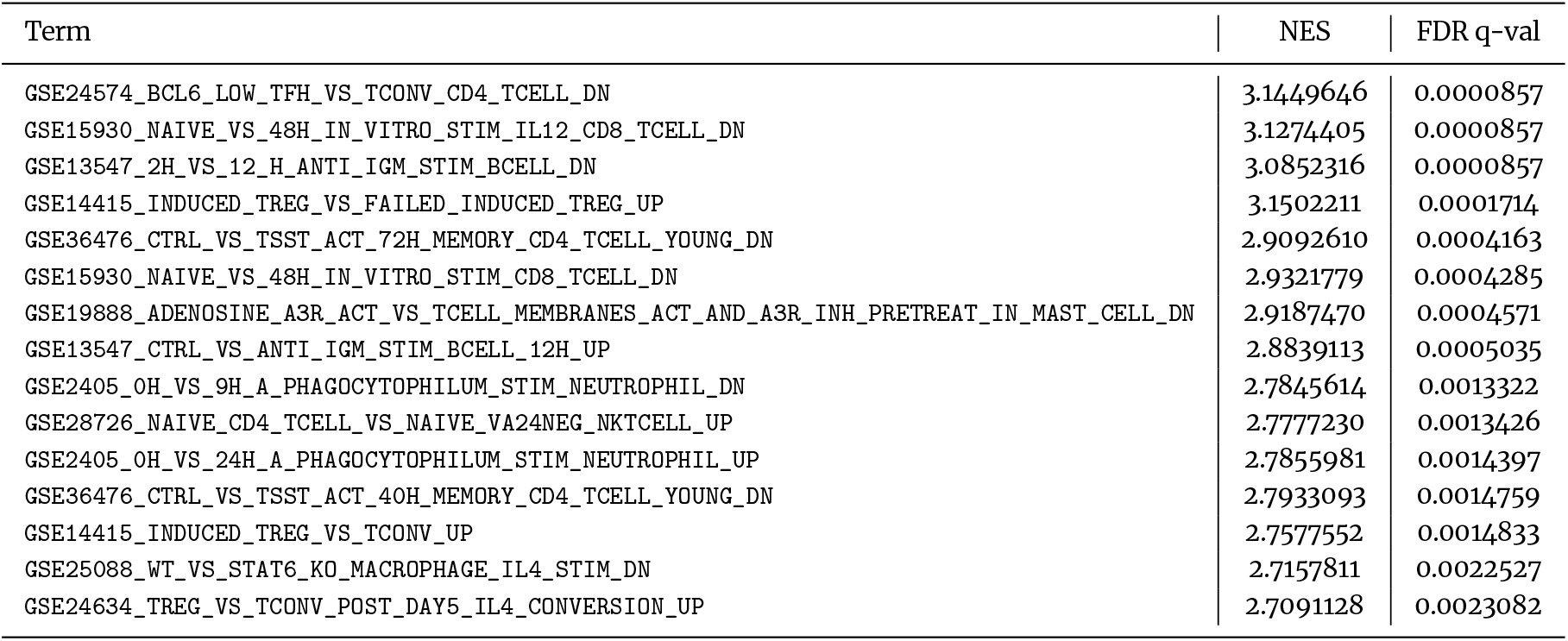
Gene set enrichment analysis against MSigDB C7 (immunlogic signature gene sets) [29, 30] for the inferred gene regulation factors across all cancer subtypes and cell types from the breast cancer use case study: Top 15 gene sets with significant enrichment (FDR *q*-value < 0.05) ranked by normalized enrichment score (NES).

**Figure S1.**
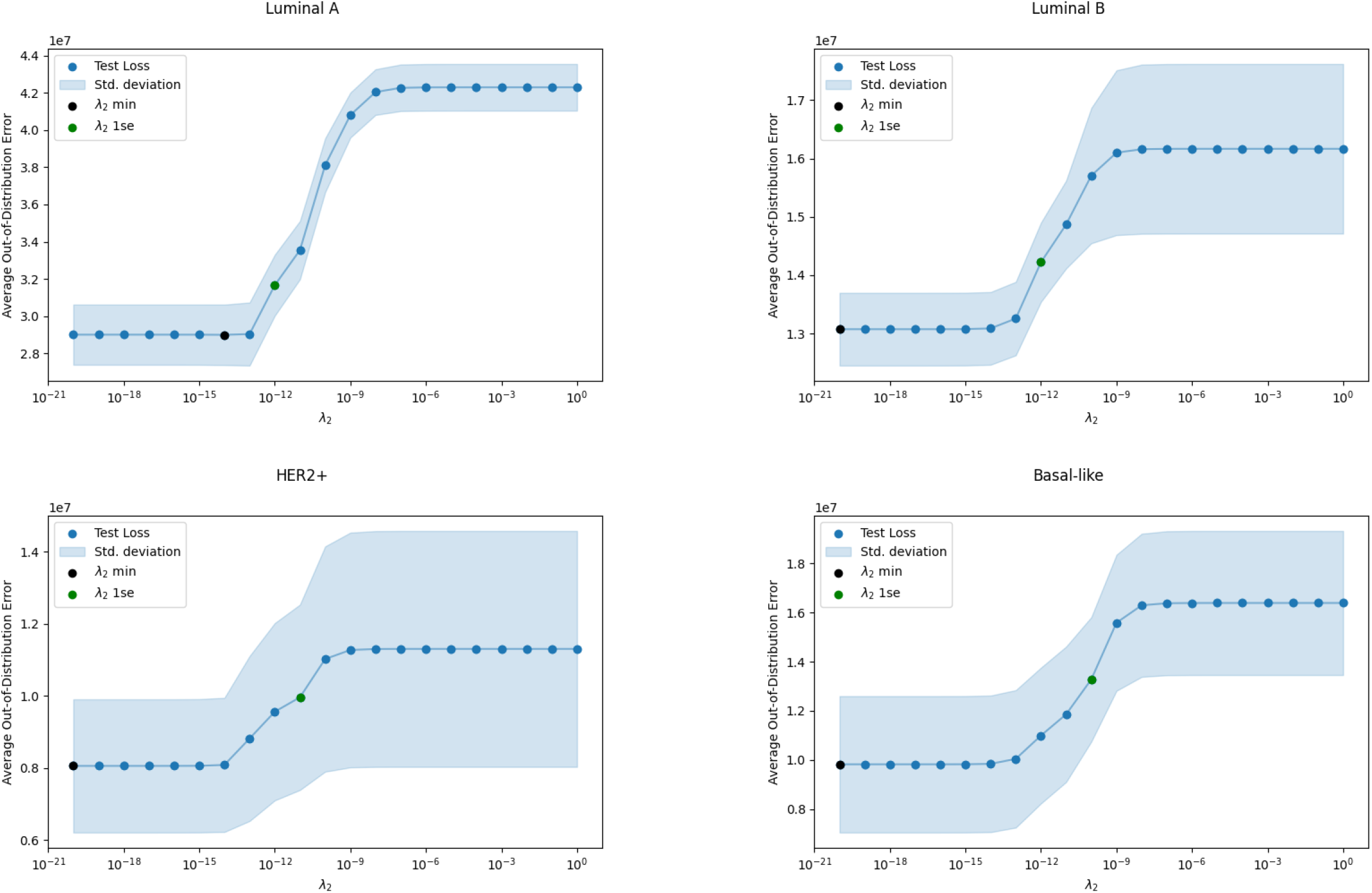
Hyperparameter search for the Deconomix+h,r model on four subsets of different breast cancer subtypes within the TCGA bulk dataset [23, 24]. Blue dots represent mean values of error on a left-out test set during five-fold cross-validation. Light blue regions represent one standard deviation around those mean values. Black dots represent the minimal λ_2_, while the green dots denote the optimal λ_2_ recommended by the one-standard-error rule (1SE rule).

**Figure S2.**
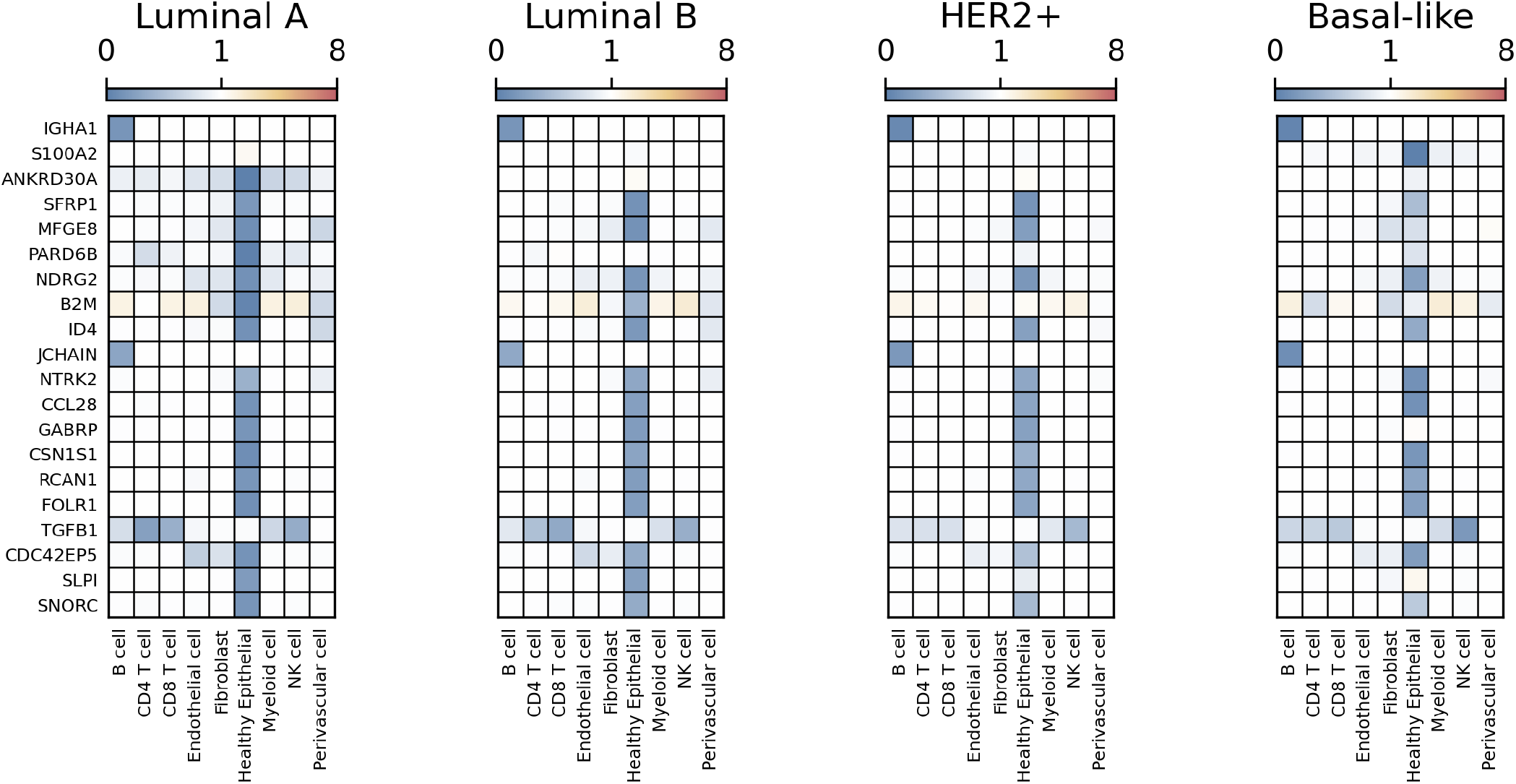
Heatmap of the 20 genes most downregulated across the four breast cancer subtypes. Values between 0 and 1 (blue) correspond to down-regulation, white to no regulation, values above 1 (yellow over orange and red) correspond to positive gene regulation.

